# 3D-Mol: A Novel Contrastive Learning Framework for Molecular Property Prediction with 3D Information

**DOI:** 10.1101/2023.08.15.553467

**Authors:** Taojie Kuang, Yiming Ren, Zhixiang Ren

## Abstract

Molecular property prediction offers an effective and efficient approach for early screening and optimization of drug candidates. Although deep learning based methods have made notable progress, most existing works still do not fully utilize 3D spatial information. This can lead to a single molecular representation representing multiple actual molecules. To address these issues, we propose a novel 3D structure-based molecular modeling method named 3D-Mol. In order to accurately represent complete spatial structure, we design a novel encoder to extract 3D features by deconstructing the molecules into three geometric graphs. In addition, we use 20M unlabeled data to pretrain our model by contrastive learning. We consider conformations with the same topological structure as positive pairs and the opposites as negative pairs, while the weight is determined by the dissimilarity between the conformations. We compare 3D-Mol with various state-of-the-art(SOTA) baselines on 7 benchmarks and demonstrate our outstanding performance in 5 benchmarks.

## 1 Introduction

Molecular property prediction can effectively and efficiently accelerate drug discovery by prioritizing promising compounds, streamlining drug development and increasing success rates. Moreover, it contributes to the comprehension of structure-activity relationships by demonstrating the influence of particular features on molecular interactions and other biological effects. Although deep learning based methods have achieved several successes in molecular property prediction, their potential is significantly constrained by the scarcity of labeled data due to the requirements of expensive and time-consuming experiments[1].

Self-supervised learning uses large amounts of unlabeled data to pretrain models to leverage unlabeled data to learn rich feature representations[2, 3, 4]. Early deep learning based methods for molecular property prediction[5, 6, 7, 8, 9] utilized NLP-based self-supervised learning methods to handle data represented by the simplified molecular input line entry system (SMILES)[10], which is a line notation with ASCII strings for the structure of chemical species. However, SMILES is not adequate for the representation of the topological structure of molecules. To address this, various pretrained molecular graph based methods, such as PretrainGNN[11], N-Gram-graph[12], MolCLR[13], and GROVER[14], have emerged. Besides, substantial methods employ graph neural network (GNN) to capture 2D information, such as MPNN[15], AttentiveFP[16], and D-MPNN[17]. Yet, these methods have ignored the 3D spatial information of molecules, which is critical as different 3D structures may lead to dissimilar molecular properties despite having the same 2D molecular topology. As an example shown in Figure 1, Thalidomide, a sedative and treatment for morning sickness in pregnant women in the 1950s, has two distinct 3D structures, R-Thalidomide and S-Thalidomide. The former has desired drug effects, while the latter has been implicated in teratogenesis. Recently, several works based on graph structures integrated with geometric information have made substantial progress in molecular property prediction tasks[18, 19, 20, 21, 22, 23, 24, 25, 22]. But they have limited 3D information extraction ability from masking-based self-supervised learning approaches and utilize only the stablest (with the lowest energy) 3D conformation while ignoring the diverse structural information from all various molecular conformations.

**Figure 1:**
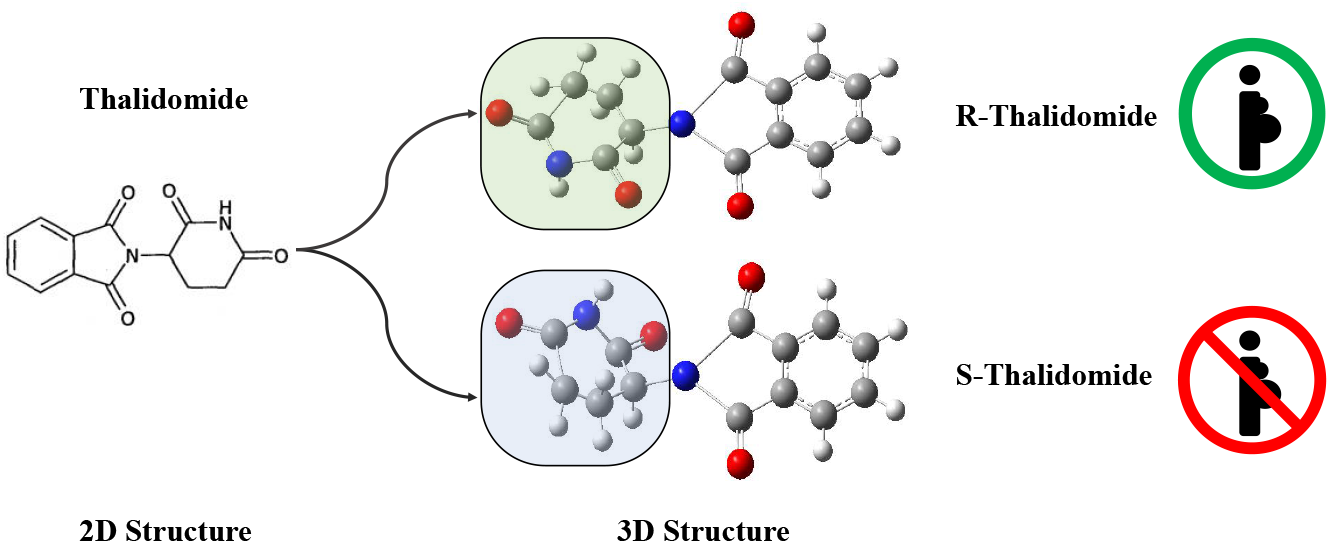
Geometric difference leads to diverse properties. Thalidomide exists in two distinct 3D stereoisomeric forms, known as R-Thalidomide and S-Thalidomide. These two molecules can be represented by the same SMILES, but they have significantly dissimilar properties. The former is recognized for its therapeutic properties, while the latter has been implicated in teratogenesis.

To address these issues, we propose a novel method, 3D-Mol, for molecular representation and property prediction. Firstly, we employ three graphs to hierarchically represent the atom-bond, bond-angle, and dihedral information of molecules. These three graphs are integrated and trained with message passing strategy as molecular embeddings. Second, using a vast amount of unlabeled data, we create a novel self-supervised method, called weighted contrastive learning, to pretrain our molecular encoder alongside the geometric approach from GearNet[26]. The proposed contrastive learning operates on various 3D molecular conformations, considering the ones from the same SMILES as positives and the opposites as negative pairs, while the weight represents the conformational similarity. The molecular encoder is then finetuned on downstream tasks to predict molecular properties. Finally, we compare our approach with several SOTA baselines on 7 molecular property prediction benchmarks[27], where our method achieves the best results on 5 benchmarks. In summary, our main contributions are as follows:

- We propose a novel molecular embedding method based on **hierarchical graph representation** to thoroughly extract the 3D spatial structural features of molecules.
- We improve the contrastive learning approach by utilizing **3D conformational information** by considering conformations with the same SMILES as positive pairs and the opposites as negative pairs, while keeping the weight to indicate the structural similarity.
- We evaluate 3D-Mol on various molecular property prediction benchmarks, showing that our model can significantly **outperform existing competitive models** on multiple tests.

## 2 Related Work

Two general methods can be used to enhance the prediction of molecular properties. One involves creating a novel molecular encoder based on molecular data for efficient latent vector extraction, and the other involves creating a unique pretraining approach to pretrain the molecular encoder using a lot of unlabeled data. The associated pieces by each are listed below.

### 2.1 Molecular Representation and Encoder

Powerful molecular representation and encoding are essential for accurate property prediction, vital in applications like molecular design and drug discovery. Proposing a novel molecular representation and encoder method is usually the first option to improve the accuracy of molecular property prediction. Some early works[21, 28, 29] learned representation from chemical fingerprints (FP), such as ECFP[21] and MACCS[30]. The others learn representation from molecular descriptors, such as SMILES. Inspired by mature NLP models, SMILES-BERT[31] used SMILES to extract molecular representations by applying the BERT[2] pretraining strategy. However, These methods depend on feature engineering, failing to capture the complete topological structure information of molecules.

Since a molecular graph is a natural representation of a molecule and conveys topological information, several research in recent years have embraced it as a means of molecular representation. GG-NN[15], DMPNN[17], and DeepAtomicCharge[32] employed a message passing strategy for molecular property prediction. AttentiveFP[16] uses a graph attention network to aggregate and update node information. The MP-GNN[33] merges specific-scale GNN and element-specific GNN, capturing various atomic interactions of multiphysical representations at different scales. MGCN[34] designed a GCN to capture multilevel quantum interactions from the conformation and spatial information of molecules.

The above mentioned works focus on 2D molecular representation, but extracting only 2D topological information from molecules is insufficient[35] and may overlook the key chemical details[36]. Recently, some research has also begun modeling 3D molecules to address this issue. SGCN[25] applies different weights according to atomic distances during the GCN-based message passing process. SchNet[18] models complex atomic interactions using Gaussian radial basis functions for potential energy surface prediction to accelerate the exploration of chemical space. DimeNet[22] proposes directional message passing to fully utilize directional information within molecules. GEM[23] develops a novel geometrically enhanced molecular representation learning method and employs a specifically designed geometric-based GNN structure. However, these methods do not fully exploit the 3D structural information of molecules and lack the ability to learn the representations of 3D conformations with the same molecular topology. To address this, we design a novel molecular representation and the corresponding encoder.

### 2.2 Self-supervised Learning on Molecules

Self-supervised learning, where BERT[2] and GPT[37] are the most prominent and successful examples, has achieved enormous success in many research fields. Inspired by these, numerous works on molecular property prediction use this approach to effectively utilize large amounts of unlabeled data for pretraining.

For one-dimensional data, SMILES is frequently used to extract molecular representations in the pretraining stage. SMILES2Vec[5] employed RNN to extract features from SMILES. ChemBERTa[9] followed RoBERTa[38] by employing masked language modeling as a pretraining task, predicting masked tokens to restore the original sentence, which helped models understand sequence semantics. SMILES Transformer[39] used a SMILES string as input to produce a temporary embedding, which is then restored to the original input by a decoder.

As the topological information of molecular graphs received greater attention, numerous pretraining methods focused on molecular graph data have been proposed. N-gram graph[12] used the n-gram method in NLP to extract representations of molecules. PretrainGNN[11] proposed a new pretraining strategy, including node-level and graph-level self-supervised pretraining tasks. GraphCL[40], MOCL[41], and MolCLR[13] performed molecular contrastive learning via GNN by proposing new molecular graph augmentation methods. MPG[42] and GROVER[14] focused on node level and graph level representation and designed the corresponding pretraining tasks for node level and graph level. iMolCLR[43], Sugar[44] and ReLMole[45] focused on the substructure of molecules, and designed the substructure pretraining task by using substructure information. However, these pretraining strategies are only targeted at learning the topological structure information of the molecule, ignoring the 3D structure information.

With the 3D structure information of molecules proven to boost molecular property prediction, recent works have focused on pretraining tasks for the 3D structure information of molecules. 3DGCN[46] introduced a relative position matrix that includes 3D positions between atoms to ensure translational invariance during convolution. GraphMVP[47] proposed a SSL method involving contrastive learning and generative learning between 3D and 2D molecular views. GEM[23] proposed a self-supervised framework using molecular geometric information by constructing a new bond angle graph, where the chemical bonds within a molecule are considered as nodes and the angle formed between two bonds is considered as the edge. Uni-Mol[48] employed the transformer to extract molecular representation by predicting atom distance. These works have not fully utilized the spatial information of molecules or the ability to learn the representation of geometric isomers. To address these issues, we use RDKit[49], an open-source toolkit for cheminformatics, to generate multiple conformations and design a weighted contrastive learning task for pretraining. We considered the ones from the same SMILES as positives and the opposites as negative pairs, while the weight represents the conformational similarity.

## 3 Method

### 3.1 Molecular Representation

We deconstruct molecular conformation into three graphs, denoted as *Mol* = {*G*_*a*−*b*_, *G*_*b*−*a*_, *G*_*d*−*a*_. Molecular raw data is represented by SMILES in most molecular databases. To extract spatial structure information from molecules, we need to use RDKit to transform the SMILES representation into 3D molecular conformations. In our method, we decompose molecular conformation into three graphs. The first graph, named atom-bond graph, is commonly used as a 2D molecular graph and is represented as *G*_*a*−*b*_ = {*V, E, P*_*atom*_, *P*_*bond*_}, where *V* is the set of atoms and *E* is the set of bonds. 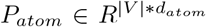, and are the attributes of atoms, and *d*_*atom*_ is the number of atom attributes. 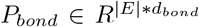, and is the attributes of bonds, and *d*_*bond*_ is the number of bond attributes. The second graph, named bond-angle graph, is represented as *G*_*b*−*a*_ = {*E, P, Ang*_*θ*_}, where *P* is a set of the plane that is comprised of 3 connected atoms. *Ang*_*θ*_ is the set of corresponding bond angles *θ*. The third graph, named the dihedral-angle graph, is represented as *G*_*d*−*a*_ = *{P, D, Ang*_*ϕ*_*}. D* represents the set of two connected planes, which connect with a bond. *Ang*_*ϕ*_ represents the corresponding dihedral angle *ϕ*. These three graphs represent an actual molecule, and help our encoder learn 3D structure information.

### 3.2 Attribute Embedding Layer

The 3D information of the molecule, such as the length of bonds and the angle between bonds, carries key chemical information. Firstly, we convert float numbers, like angle and bond length, to latent vectors. Referring to the previous work[24], we employed several RBF layers to encode different geometric factors:

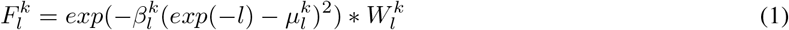

where 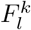 is the k-dimensional feature of bond length *l*, and 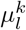 and 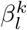 are the center and width of *l* respectively. 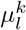 is 0.1*k* and 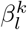 is 10. Similarly, the k-dimensional features of 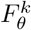 and 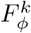 of x are computed as:

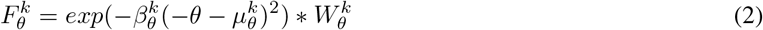

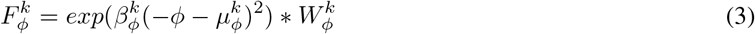

where 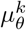 and 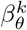 are the center and width of *θ*, and 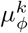 and 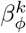 are the center and width of *ϕ*. The centers of bond angle and dihedral angle are represented as 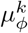 and 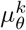, and the numerical values of them are *π*/K, where K is the number of feature dimensions.

For the other attributes of atom and bond, we represent them with *P*_*atom*_ and *P*_*bond*_. Inspired by NLP, we embed them with the word embedding function. The initial features of atoms and bonds are represented as 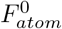 and 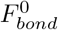 respectively.

### 3.3 3D-Mol Layer

Inspired by GEM[23], we employ message passing strategy to send and receive messages in 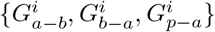 For the *i*_*th*_ layer in 3D-Mol, the information of 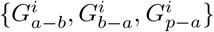 will be updated by message passing neural network. The message passing of the latter graph needs the information of the former graph. The overview is shown in figure 3, and the details are as follows:

**Figure 2:**
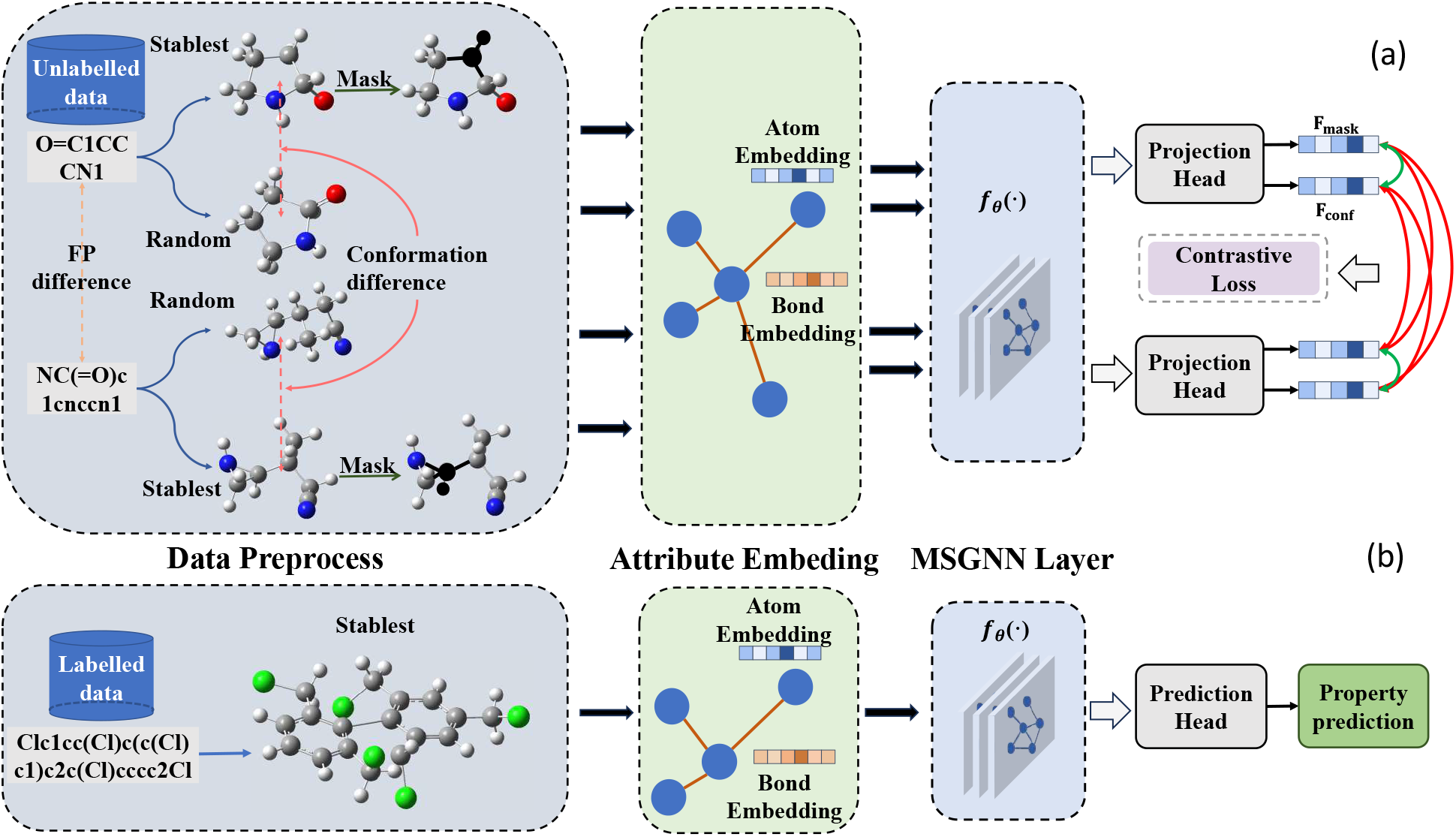
The overview of the 3D-Mol model framework. a) In the pretraining stage, we employ weighted contrastive learning to effectively pretrain our model. In addition to using the mask strategy for graph data augmentation, we consider conformations from the same SMILES as positive pairs, while the weight represents the conformational similarity. Conversely, distinct topological structures are treated as negative pairs, and the weight is dependent on FP differences. b) In the finetuning stage, we refine the well-pretrained encoder using diverse downstream datasets, followed by supervised learning.

**Figure 3:**
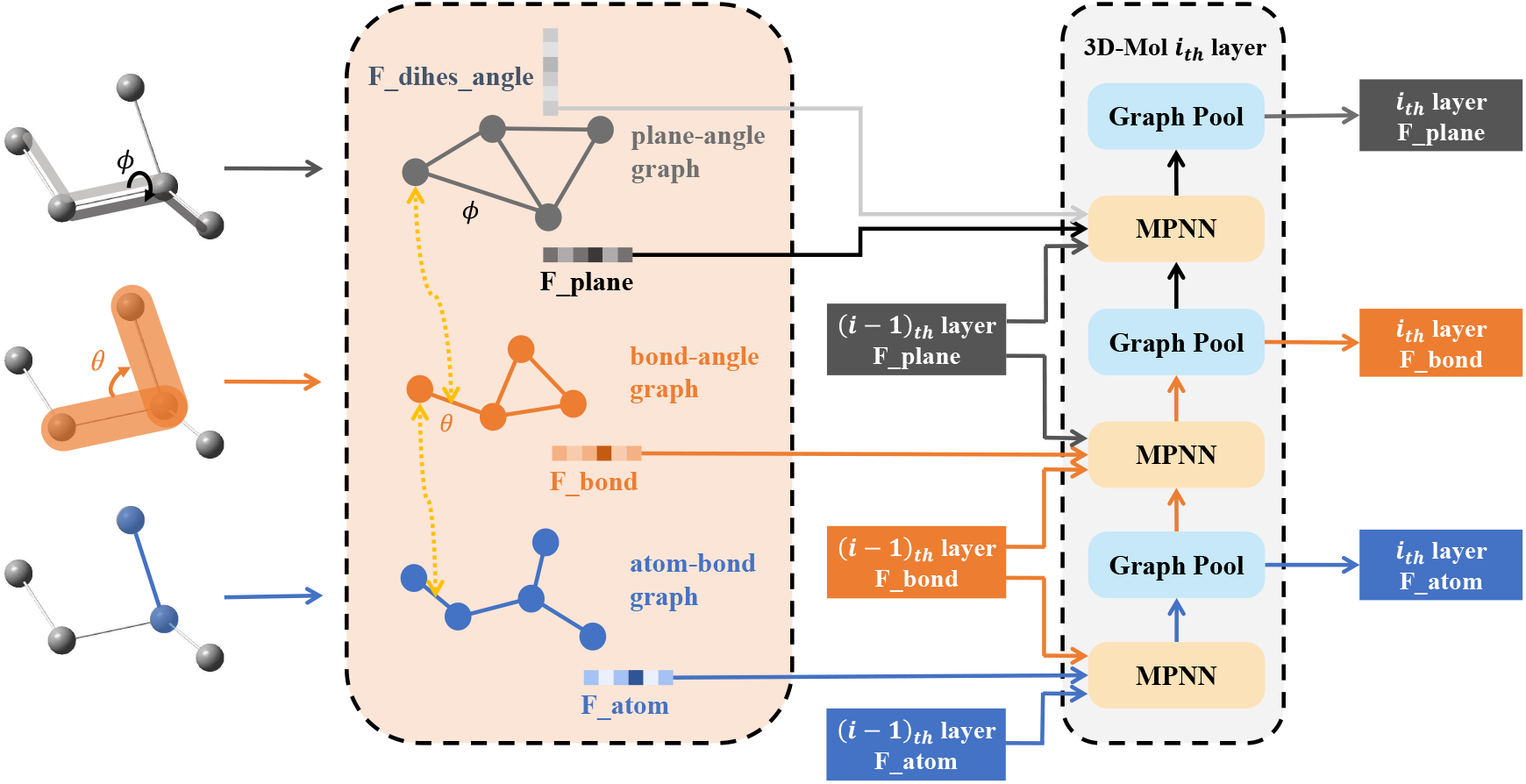
The overview of the 3D-Mol encoder layer. The 3D-Mol encoder layer comprises three steps. Firstly, employing a message passing strategy, nodes in each graph exchange messages with their connected edges, leading to the updating of edge and node latent vectors. Secondly, the edge latent vector from the lower-level graph is transmitted to the higher-level graph as part of the node latent vector. Finally, the iteration is performed n times to derive the *n*_*th*_ node latent vector, from which we extract the molecular latent vectors.

First, we use 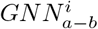 to aggregate the message and update the atom and bond latent vectors in 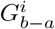. Given an atom v, its representation vector 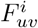 is formalized by:

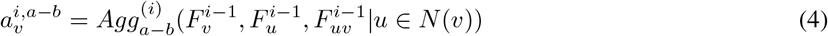

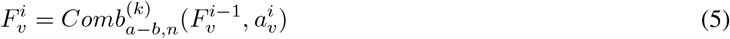

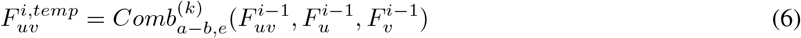

where *N* (*v*) is the set of neighbors of atom v in 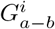, and 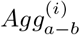 is the aggregation function for aggregating messages from the atom neighborhood. 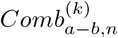 and 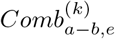 are the update functions for updating the latent vectors of atom and bond, respectively. 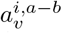 is the information from the neighboring atom and the corresponding bond after being aggregated. 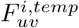 is the temporary bond latent vectors of bond *uv* in *i*_*th*_ layer and is part of the bond latent vectors in 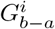. Massage passing in 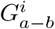 updates the latent vectors of atoms and bonds and helps the model learn the topology information of the molecule.

Then, we use 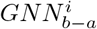 to aggregate the message and update the bond and plane vectors in 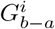. Given a bond *uv*, its latent vector 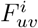 is formalized by:

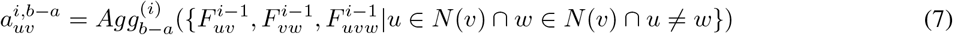

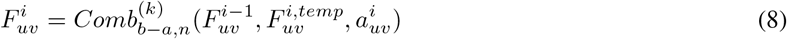

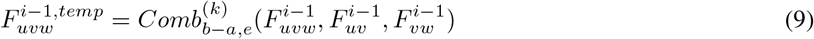

where 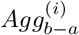 is the aggregation function for aggregating messages from the bond neighborhood. 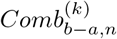 and 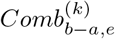 are the update functions for updating the bond and plane latent vectors. 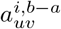 is the information from the neighboring bond and the corresponding bond angle after being aggregated. 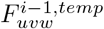 is the temporary plane latent vectors of plane *uvw* in *i*_*th*_ layer and is part of the plane latent vectors in 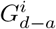.

After processing the 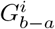, we use 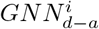 to aggregate the message and update the plane latent vector in 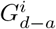 Given a plane which is constructed by nodes u, v, w and bonds *uv, vw*, its latent vector 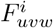 is formalized by:

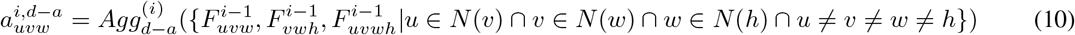

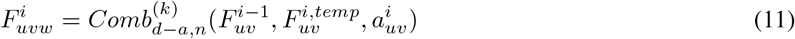

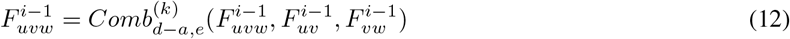

where 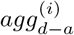 is the aggregation function for aggregating messages from the plane neighborhood. 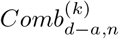 and 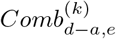 are the update functions for updating the plane latent vector. 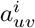 is the information from the neighboring plane and the corresponding dihedral angle after being aggregated. Massage passing in 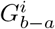 and 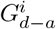 updates the latent vectors of bonds and planes and helps the model learn the 3D structure information of the molecule.

The representation vectors of the atoms at the final iteration are integrated to gain the molecular representation vector *F*_*mol*_ by the Readout function, which is formalized as:

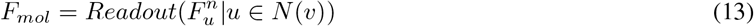

where *F*^*n*^ is the last 3D-Mol layer output. The molecular latent vector *F*_*mol*_ is used to predict molecular properties.

#### 3.3.1 Geometry task

Since geometry information has been demonstrated to be important for molecular property prediction[22], we also employ geometry tasks as the pretraining method. For bond angle and dihedral angle prediction, we sample adjacent atoms to better capture local structural information. Since angular values are more sensitive to errors in protein structures than distances, we use discretized values for prediction. The following are the loss functions for the local geometry task:

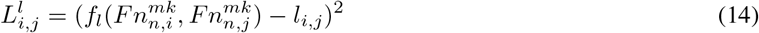

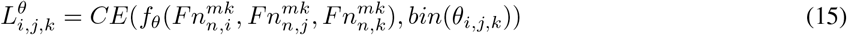

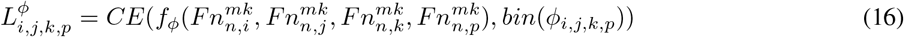

where *f*_*ϕ*_(.), *f*_*θ*_(.) and *f*_*l*_ are the MLPs for the local geometry task, and 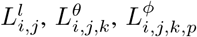 and 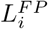 are the loss functions for each task. *CE*(.) is the cross entropy loss, and *bin*() is used to discretize the bond angle and dihedral angle. 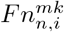 is the latent vector of node i after masking the corresponding sampled items in each task.

In addition to the aforementioned pretraining tasks to capture global molecular information, we leverage masked molecular latent vectors for FP prediction and atom distance prediction, effectively incorporating latent representations to enrich the predictive capability. The following are the loss functions for the global geometry task:

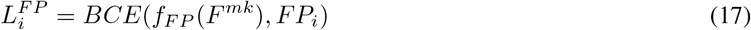

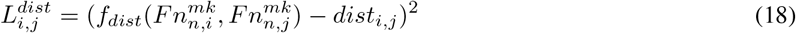

where *f*_*F P*_ and *f*_*dist*_(.) are the MLPs for global geometric tasks, and 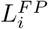 and 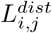 are the loss functions for each task. *BCE*(.) is binary cross entropy loss. *F*^*mk*^ is the latent vector of the masking molecule.

## 4 Experiment

In this section, we conduct experiments on 7 benchmark datasets in MoleculeNet to demonstrate the effectiveness of 3D-Mol for molecular property prediction. We use a large amount of unlabeled data and our proposed contrastive learning strategy to pretrain the 3D-Mol, and then we use the downstream task to finetune the well-pretrained model and predict the molecular property. We compare it with a variety of SOTA methods and conduct several ablation studies to confirm the effectiveness of 3D-Mol encoder and our pretraining strategy.

### 4.1 Datasets and Setup

#### 4.1.1 Pretraining stage

We use 20 million unlabeled molecules to pretrain 3D-Mol. The unlabeled data is extracted from ZINC20 and PubChem, both of which are publicly accessible databases containing drug-like compounds. To ensure consistency with prior research[23], we randomly selected 90% of these molecules for training purposes, while the remaining % were set aside for evaluation. The raw data obtained from ZINC20 and PubChem was provided in SMILES format. In order to convert the SMILES representations into molecular conformations, we employed RDKit and applied the ETKDG method. For our model, we use the Adam optimizer with a learning rate of 1e-3. The batch size is set to 256 for pretraining and 32 for finetuning. The hidden size of all models is 256. The geometric embedding dimension K is 64, and the number of angle domains is 8. The hyperparameters *λ*_*conf*_ and *λ*_*fp*_ are both set to 0.5.

#### 4.1.2 Finetuning stage

We use 7 molecular datasets obtained from MoleculeNet to demonstrate the effectiveness of 3D-Mol. These datasets encompass a range of biophysics datasets such as BACE, physical chemistry datasets like ESOL, and physiology datasets like Tox21. In the fine-tuning stage, we employed nine molecular datasets obtained from MoleculeNet. These datasets encompass a range of biophysics datasets such as BACE, physical chemistry datasets like ESOL, and physiology datasets like Tox21. Table 1 provides a summary of the statistical information for these datasets, while the remaining details are outlined below:

**Table 1:** Statistics information of datasets.

| Dataset | # Tasks | Task Type | # Molecules |
| --- | --- | --- | --- |
| BACE | 1 | Classification | 1513 |
| Sider | 27 | Classification | 1427 |
| Tox21 | 12 | Classification | 7831 |
| ToxCast | 617 | Classification | 8597 |
| ESOL | 1 | Regression | 1128 |
| FreeSolv | 1 | Regression | 643 |
| Lipophilicity | 1 | Regression | 4200 |

- BACE. The BACE dataset provides both quantitative (IC50) and qualitative (binary label) binding results for a set of inhibitors targeting human *β*-secretase 1 (BACE-1).
- Tox21. The Tox21 initiative aims to advance toxicology practices in the 21st century and has created a public database containing qualitative toxicity measurements for 12 biological targets, including nuclear receptors and stress response pathways.
- Toxcast. ToxCast, an initiative related to Tox21, offers a comprehensive collection of toxicology data obtained through in vitro high-throughput screening. It includes information from over 600 experiments and covers a large library of compounds.
- SIDER. The SIDER database is a compilation of marketed drugs and their associated adverse drug reactions (ADRs), categorized into 27 system organ classes.
- ESOL. The ESOL dataset is a smaller collection of water solubility data, specifically providing information on the log solubility in mols per liter for common organic small molecules.
- FreeSolv. The FreeSolv database offers experimental and calculated hydration-free energy values for small molecules dissolved in water.
- Lipo. Lipophilicity is a crucial characteristic of drug molecules that affects their membrane permeability and solubility. The Lipo dataset contains experimental data on the octanol/water distribution coefficient (logD at pH 7.4).

Following the previous work[23], we partitioned the datasets into train/validation/test sets in an 80/10/10 ratio for downstream tasks, and we use scaffold splitting and report the mean and standard deviation by the results of 3 random seeds.

### 4.2 Metric

Consistent with prior studies, we adopt the average ROC-AUC as the evaluation metric for the classification datasets (BACE, SIDER, Tox21 and ToxCast), which is a widely used metric for assessing the performance of binary classification tasks. For the regression datasets (ESOL, FreeSolv and Lipophilicity), we utilize the RMSE as the evaluation metric.

### 4.3 Result

a. **To validate the efficacy of our proposed method, we compare it with several baseline methods**. The baseline methods are as follows: N-Gram[12] generates a graph representation by constructing node embeddings based on short walks. PretrainGNN[11] implements several types of self-supervised learning tasks. 3D Infomax[50] maximizes the mutual information between learned 3D summary vectors and the representations of a GNN. MolCLR[13] is a 2D-2D view contrastive learning model that involves atom masking, bond deletion, and subgraph removal. GraphMVP[47] used 2D-3D view contrastive learning approaches. GROVER[14] focused on node level and graph level representation and corresponding pretraining tasks for node level and graph level. GEM[23] employs predictive geometry self-supervised learning schemes that leverage 3D molecular information. Uni-Mol[48] enlarges the application scope and representation ability of molecular representation learning by using transformer. As shown in Table 2, Our method gets the best result in 5 datasets and the second best result in 1 dataset. Furthermore, our method achieves overwhelming performance on BACE by a large margin. That shows that our method is better at extracting molecular information.
b. **To validate the efficacy of our proposed model 3D-Mol encoder, we compare it with several baseline molecular encoder without pretraining**. The baseline molecular encoders are as follows: DMPNN[17] employed a message passing scheme for molecular property prediction. AttentiveFP[16] is an attention-based GNN that incorporates graph-level information. MGCN[34] designed a hierarchical GNN to directly extract features from conformation and spatial information, followed by multilevel interactions. HMGNN[24] leverages global molecular representations through an attention mechanism. SGCN[25] applies different weights according to atomic distances during the GCN-based message passing process. DimeNet[22] proposes directional message passing to fully utilize directional information within molecules. GEM[23] employed message passing strategy to extract 3D molecular information. We present the experimental result to show the efficiency of our 3D-Mol model. As the result shown in Table 3, 3D-Mol encoder significantly outperforms all the baselines on both types of tasks and improves the performance over the best baselines with 2% and 11% for classification and regression tasks, respectively, since 3D-Mol incorporates geometrical parameters.
c. **To validate the efficacy of our proposed pretraining task, we study the performance of 3D-Mol encoder in three scenarios: without pretraining, pretraining by geometrical tasks, and pretraining by geometrical tasks with weighted contrastive learning**. The result in Table 4 shows that geometry tasks significantly improve the performance of 3DGNN. Compared with the pretraining method combined with weighted contrastive learning and the method without pretraining, The former is better in all datasets. Compared with the pretraining method combined with weighted contrastive learning and the pretraining method without weighted contrastive learning, The former is better in all datasets but SIDER, Tox21, and FreeSolv. In general, our pretraining method is better for improving the 3DGNN performance since model learn the 3D structure information in the pretraining stage.

**Table 2:** Benchmarking the 3D-Mol and other pretraining methods. We compare the performance on the 7 molecular property prediction tasks, marking the best results in bold and underlining the second best.

| model | Classification (ROC-AUC %, higher is better ↑) | | | | Regression (RMSE, lower is better ↓) | | |
| --- | --- | --- | --- | --- | --- | --- | --- |
|  | BACE | SIDER | Tox21 | ToxCast | ESOL | FreeSolv | Lipophilicity |
| N-Gram<sub>RF</sub> | 0.779<sub>0.015</sub> | 0.668<sub>0.007</sub> | 0.743<sub>0.004</sub> | — | 1.074<sub>0.107</sub> | 2.688<sub>0.085</sub> | 0.812<sub>0.028</sub> |
| N-Gram<sub>XGB</sub> | 0.791<sub>0.013</sub> | 0.655<sub>0.007</sub> | 0.758<sub>0.009</sub> | — | 1.083<sub>0.107</sub> | 5.061<sub>0.744</sub> | 2.072<sub>0.030</sub> |
| PretrainGNN | 0.845<sub>0.007</sub> | 0.627<sub>0.008</sub> | 0.781<sub>0.006</sub> | 0.657<sub>0.006</sub> | 1.100<sub>0.006</sub> | 2.764<sub>0.002</sub> | 0.739<sub>0.003</sub> |
| 3DInfomax | 0.797<sub>0.015</sub> | 0.606<sub>0.008</sub> | 0.644<sub>0.011</sub> | 0.745<sub>0.007</sub> | 0.894<sub>0.028</sub> | 2.337<sub>0.107</sub> | 0.695<sub>0.012</sub> |
| GraphMVP | 0.812<sub>0.009</sub> | 0.639<sub>0.012</sub> | 0.759<sub>0.005</sub> | 0.631<sub>0.004</sub> | 1.029<sub>0.033</sub> | — | 0.681<sub>0.010</sub> |
| GROVER<sub>base</sub> | 0.826<sub>0.007</sub> | 0.648<sub>0.006</sub> | 0.743<sub>0.001</sub> | 0.654<sub>0.004</sub> | 0.983<sub>0.090</sub> | 2.176<sub>0.052</sub> | 0.817<sub>0.008</sub> |
| GROVER<sub>large</sub> | 0.810<sub>0.014</sub> | 0.654<sub>0.001</sub> | 0.735<sub>0.001</sub> | 0.653<sub>0.005</sub> | 0.895<sub>0.017</sub> | 2.272<sub>0.051</sub> | 0.823<sub>0.010</sub> |
| MolCLR | 0.824<sub>0.009</sub> | 0.589<sub>0.014</sub> | 0.750<sub>0.002</sub> | — | 1.271<sub>0.040</sub> | 2.594<sub>0.249</sub> | 0.691<sub>0.004</sub> |
| GEM | 0.856<sub>0.011</sub> | **0.672**<sub>0.004</sub> | 0.781<sub>0.005</sub> | 0.692<sub>0.004</sub> | 0.798<sub>0.029</sub> | 1.877<sub>0.094</sub> | 0.660<sub>0.008</sub> |
| Uni-Mol | <u>0.857</u><sub>0.005</sub> | <u>0.659</u><sub>0.013</sub> | **0.796**<sub>0.006</sub> | <u>0.696</u><sub>0.001</sub> | <u>0.788</u><sub>0.029</sub> | <u>1.620</u><sub>0.035</sub> | <u>0.603</u><sub>0.010</sub> |
| 3D-Mol | **0.875**<sub>0.004</sub> | 0.656<sub>0.002</sub> | <u>0.786</u><sub>0.003</sub> | **0.697**<sub>0.003</sub> | **0.783**<sub>0.009</sub> | **1.617**<sub>0.050</sub> | **0.598**<sub>0.018</sub> |

**Table 3:** Benchmarking the 3D-Mol and other non-pretraining methods. We compare the performance on the 7 molecular property prediction tasks, marking the best results in bold and underlining the second best.

| model | Classification (ROC-AUC %, higher is better ↑) | | | | Regression (RMSE, lower is better ↓) | | |
| --- | --- | --- | --- | --- | --- | --- | --- |
|  | BACE | SIDER | Tox21 | ToxCast | ESOL | FreeSolv | Lipophilicity |
| DMPNN | 0.809<sub>0.006</sub> | 0.570<sub>0.007</sub> | 0.759<sub>0.007</sub> | 0.655<sub>0.3</sub> | 1.050<sub>0.008</sub> | 2.082<sub>0.082</sub> | 0.683<sub>0.016</sub> |
| AttentiveFP | 0.784<sub>0.000</sub> | 0.606<sub>0.032</sub> | 0.761<sub>0.005</sub> | 0.637<sub>0.002</sub> | 0.877<sub>0.029</sub> | 2.073<sub>0.183</sub> | 0.721<sub>0.001</sub> |
| MGCN | 0.734<sub>0.008</sub> | 0.587<sub>0.019</sub> | 0.741<sub>0.006</sub> | — | — | — | — |
| SGCN | — | 0.559<sub>0.005</sub> | 0.766<sub>0.002</sub> | 0.657<sub>0.003</sub> | 1.629<sub>0.001</sub> | 2.363<sub>0.050</sub> | 1.021<sub>0.013</sub> |
| HMGNN | — | <u>0.615</u><sub>0.005</sub> | 0.768<sub>0.002</sub> | 0.672<sub>0.001</sub> | 1.390<sub>0.073</sub> | 2.123<sub>0.179</sub> | 2.116<sub>0.473</sub> |
| DimeNet | — | 0.612<sub>0.004</sub> | <u>0.774</u><sub>0.006</sub> | 0.637<sub>0.004</sub> | 0.878<sub>0.023</sub> | 2.094<sub>0.118</sub> | 0.727<sub>0.019</sub> |
| GEM | <u>0.828</u><sub>0.012</sub> | 0.606<sub>0.010</sub> | 0.773<sub>0.007</sub> | <u>0.675</u><sub>0.005</sub> | 0.832<sub>0.010</sub> | <u>1.857</u><sub>0.071</sub> | <u>0.666</u><sub>0.015</sub> |
| 3D-Mol<sub>w.o pre</sub> | **0.832**<sub>0.005</sub> | **0.624**<sub>0.013</sub> | **0.780**<sub>0.004</sub> | **0.682**<sub>0.007</sub> | **0.794**<sub>0.027</sub> | **1.769**<sub>0.039</sub> | **0.663**<sub>0.004</sub> |

**Table 4:** Ablation study. We study the performance of 3D-Mol encoder in three scenarios: without pretraining, pretraining by geometrical tasks, and pretraining by geometrical tasks with weighted contrastive learning. We then mark the best results in bold and underline the second best.

| model | Classification (ROC-AUC %, higher is better ↑) | | | | Regression (RMSE, lower is better ↓) | | |
| --- | --- | --- | --- | --- | --- | --- | --- |
|  | BACE | SIDER | Tox21 | ToxCast | ESOL | FreeSolv | Lipophilicity |
| 3DGNN | **0.875**<sub>0.004</sub> | <u>0.656</u><sub>0.002</sub> | <u>0.786</u><sub>0.003</sub> | **0.697**<sub>0.003</sub> | **0.783**<sub>0.009</sub> | <u>1.617</u><sub>0.050</sub> | **0.598**<sub>0.018</sub> |
| 3DGNN<sub>wo.pre</sub> | 0.832<sub>0.005</sub> | 0.624<sub>0.013</sub> | 0.780<sub>0.004</sub> | 0.682<sub>0.007</sub> | <u>0.794</u><sub>0.027</sub> | 1.769<sub>0.039</sub> | 0.674<sub>0.007</sub> |
| 3DGNN<sub>wo.clweighted</sub> | <u>0.874</u><sub>0.006</sub> | **0.661**<sub>0.005</sub> | **0.790**<sub>0.003</sub> | <u>0.693</u><sub>0.005</sub> | 0.795<sub>0.014</sub> | **1.557**<sub>0.003</sub> | <u>0.607</u><sub>0.006</sub> |

## 5 Conclusion

In this paper, we propose a novel molecular model framework, named 3D-Mol, to predict molecular properties. We propose a novel hierarchical graph-based molecular embedding for extracting full 3D structural information, enhancing contrastive learning with positive and negative pairs based on 3D conformational data to indicate structural similarity, and our 3D-Mol model outperforms 7 benchmarks in molecular property prediction. Our method can effectively empower molecular design and screening, revealing more potentials for future artificial intelligence aided drug discovery.

## Acknowledents

The research was supported by the PengCheng Laboratory and by PengCheng Laboratory Cloud-Brain.

